# Improving the visualization of viruses in soil

**DOI:** 10.1101/2024.09.30.615710

**Authors:** Amar D. Parvate, Trinidad Alfaro, Regan McDearis, Amy Zimmerman, Kirsten Hofmockel, William C. Nelson, James E. Evans

**Author notes:** Corresponding Author: James E. Evans.

## Abstract

Viruses are numerically the most abundant forms on Earth, and most are present in soil. Even though viruses are highly abundant in soil and critical to rhizosphere function, visualizing the diverse morphotypes within soil has been challenging. The difficulty is primarily due to the heterogenous nature of isolated suspensions that typically contain nanometer to micron scale debris which renders protein crystallography for structural studies unfeasible and hinders cryo-electron microscopy due to ice thickness and contrast issues. Here we employed and compared a simple spin filtration method to cleanup solutions of extracted viruses for direct observation with cryo-electron microscopy. Although relatively simple, the method employs common physical biochemical separation steps to remove large and small debris which dramatically improves image quality and preservation of structural features to permit visualizing morphotypes not typically seen with conventional negative stain approaches. In addition to tailed and non-tailed polyhedral phages, several under reported or novel morphotypes of soil viruses are directly visualized as a particle library with both 2D and 3D information.

## INTRODUCTION

Viruses are numerically the most abundant forms on Earth with estimates of up to 10^31^ total virus particles worldwide [1, 2] and are being explored as alternatives to pesticides, antibiotics or as source of genes encoding molecules of biotechnological importance for commercialization purposes [3-5]. Interestingly, the majority of viruses are believed to be present in soil where they are estimated to range between 10^5^ to 10^9^/ g and outnumber bacteria by 5-1,000 fold [6]. Viruses are known to integrate themselves into complex soil ecosystems and interact with microbial communities [7]. In the case of agricultural land, multiple factors impact the soil virome including the use of fertilizers, the type of fertilizer, as well as type of crops grown [8]. Furthermore, the soil type, moisture content, season of sampling, and depth of sampling can all impact the number and types of viruses observed from sample to sample [9]. This has largely limited scientific study to viruses or phages which are known to either positively or negatively impact human activities, livestock, or crops, and to a limited number of viruses which have been successfully isolated and propagated through a host [10, 11]. As of 2023, soil viromes constituted <1% of the total viromes available in public databases, making them some of the least studied forms of life on the planet [12].

“Soil” is a deceptively simple term applied to an extremely complex layer of matter covering the entire planet which harbors its own ecosystems consisting of everything from viruses to unicellular microbes, fungi as well as the rhizosphere. Soil contains solid, liquid and gaseous phases with stoichiometries varying over time and space. For soil virome studies, typically soil in the top few inches is utilized [2]. Viruses regulate microbial communities in soil environments in cycles, seasonally and spatiotemporally and may influence soil nutrient cycles in lands under cultivation [2, 13]. Negative stain TEM is the dominant approach used in already documented studies highlighting the morphological diversity of soil viruses with nanoscale imaging. Major benefits for negative stain TEM are that it is cheap and easy to perform, the resulting heavy metal stained grids are stable at room temperature for years and can be reimaged multiple times, and in general this method requires ∼10-100x less sample than other approaches like cryo-electron microscopy. One of the most comprehensive examples of high-quality negative stain TEM analysis of the soil virome was reported recently and focused on the discovery and characterization of giant virus-like particles in forest soil[14]. While they presented stunning images, the authors also acknowledged that their method of negative staining with heavy metal and high salt content in conjunction with a chemical fixation step and dehydration could perturb the viral ultrastructure resulting in shrinking and other artifacts. They also mention that “it is not possible to prove the viral nature of any particles analyzed only by negative staining TEM, regardless of how similar to published virus structures they may look,” which is partly due to the inability of negative staining to capture the full structural details and whole 3D structure of large particles. This is a widely recognized limitation of negative staining which generally fails to preserve high resolution features beyond ∼16 Angstrom for non-crystalline samples, fails to stain sugar or glycan surfaces, and fails to capture the full height and context of the virus components depending on stain depth and local area reproducibility. Several researchers have also reported that the method of extraction influences the virus morphotypes observed using negative stain electron microscopy [6]. The difference in extraction methods includes but is not limited to factors like buffer composition, type of soil, and whether a de-clumping step has been utilized, as these can interfere with the uniformity of the negative stain.

Due to these historic challenges for direct morphological analysis of soil viromes in native-like and hydrated (not chemically fixed or dehydrated) conditions, many studies are currently exploring the soil virome with only genomic methods[15, 16]. However, both genomic and morphological characterization remain critical aspects for taxonomic classification of viral Families, and both are needed to fully understand the diversity of viruses present in soil [17]. Cryo-electron microscopy (cryo-EM) is a well-established technique for imaging biological samples in near-native suspension conditions (i.e. monodisperse suspensions without any use of fixatives or heavy metal salt stains) and allows the preservation of atomic details while permitting direct visualization of material up to 0.8-1 micrometer in thickness which is an ideal range for most virus particles. Here we present a sample preparation approach that quickly and simply converts native soil isolates into a suspension compatible with high-resolution cryo-EM workflows for visualizing the soil virome with 2D and 3D analyses. Using this workflow, we observed several different morphotypes of tail-less and tailed phages with symmetrical heads and tail components, as well as a diverse array of other morphotypes including round, elongated and polymorphic viruses, and some morphotypes not previously reported using direct visualization.

## METHODS

### Field Site Description

Soil was collected on 7 March 2023 from the Tall Wheatgrass Irrigation Field Trial at the Washington State University Irrigated Agriculture Research and Extension Center (WSU IAREC) in Prosser, WA. As described previously [18], the site has aridisol soils with low organic matter content and alkaline pH and was planted with Tall Wheatgrass in 2018. Irrigation treatments from 25% to 100% Field Water Capacity (FWC) have been ongoing since spring 2019. Soil was collected from a random location within plot 23, which is planted with the Alkar cultivar of Tall Wheatgrass and irrigated at 25% FWC, using a 5 cm diameter hammer corer. The soil from 5-15 cm depth was used in this study. Soil was transported to the laboratory on blue ice, where it was homogenized and sieved (<2 mm) prior to further sample processing.

### Enrichment of virus-like particles from soil

Virus-sized particles were enriched from freshly collected soils (maintained at 4ºC after homogenization) following methods adapted from Santos-Medellín et al. [13]. For each sample, 20 g of soil was weighed into 50 mL tubes. Each tube received 18 mL of protein supplemented phosphate-buffered saline solution (PPBS: 2% bovine serum albumin, 10% phosphate-buffered saline, 1% potassium citrate, and 150 mM MgSO4) and was mixed at 300 rpm for 10 minutes at 4°C. After mixing, tubes were centrifuged at 4,000 x g for 10 minutes at 4°C. The supernatant was collected, and the soil pellet was resuspended in 18 mL PPBS, mixed as above, and centrifuged. These steps were repeated once more. The pooled supernatant was then centrifuged twice at 10,000 x g for 8 minutes at 4°C for removal of remaining soil particles. The supernatant was then filtered through a 0.22 μm filter for removal of bacterial cells. To concentrate the virus-like particles (VLP), the filtrate was ultracentrifuged in a 45 Ti rotor at 32,000 x g for 3 hours at 4°C, and the supernatant discarded. To resuspend the VLP pellet, 300 µL of cell culture grade water was added directly to the surface of the pellet, incubated at 4°C for 1 hour at 150 rpm, and gently resuspended via pipetting.

### Clean-up step before TEM imaging

The resulting VLP suspensions were subsampled (40 µL) for direct visualization by cryo-EM. Each suspension was diluted ∼9-10X in TBD buffer (350-400 µL) and loaded on a Pleuriselect 1 µm membrane filter (Cat # SKU 46-10001-15) to separate large micron size debris. Samples were centrifuged and the filtrate was loaded on a 300 kDa MWCO Amicon spin concentrator (Millipore) to further concentrate the retentate to ∼30 µL by centrifugation. The filtrate was then loaded on a 100 kDa MWCO spin concentrator and centrifuged. The retentate from this step was concentrated to 30 µL. Alternatively, the filtrate from the Pleuriselect filter was loaded directly onto a 100 kDa MWCO filter and the retentate was concentrated to 30 µL. All centrifugation steps were performed at 5000-10,000 x g at 4°C for 15 min.

### Grid preparation for TEM imaging

Aliquots of soil samples before the Pleuriselect step, and retentates from the 300 and 100 kDa MWCO filters were used for preparing cryo-EM grids. Briefly, 3 µL of sample was loaded onto glow discharged holey carbon grids (Quantifoil, 300 mesh, R1/2). Samples were blotted for 3s and plunge frozen in liquid ethane using a Leica EM GP2 to obtain vitrified ice for imaging. For negative staining, 2 µL suspension from the 300 kDa MWCO retentate was loaded on holey carbon grids and excess sample was blotted away. Staining was performed using Nano-W™ (Ted Pella Inc.), following conventional procedures of negative stained sample preparation [18]

### TEM, cryo-EM and cryo-ET

Samples were loaded on a 300 keV (Thermo Fisher Scientific) Titan Krios and all screening and data collection was performed using standard EPU software. Images were collected either at nominal magnifications ranging from 135x, 580x, 11500x for low dose screening and 42000x for data collection using a K3 direct electron detector (Gatan Inc.) fitted with a Bioquantum energy filter and a slit width of 20 eV. Dose was not optimized for low dose screening images, but exposure was kept to 1 s. For data collection, movies were collected at a pixel size of 1.1 Å in counting mode, using a defocus of -1 to -3 µm, at an exposure of 2-2.5 s exposure across 32 frames resulting in a total dose of 25-28 e^-^/Å^2^/s. Motion correction was performed using standard parameters in CryoSparc v4.2 and micrographs were Fourier cropped to 0.25 x [19]. Motion corrected micrographs were inspected and sorted manually in Fiji [20] to visualize different virus morphotypes. For cryo-ET, tilt series were collected from +/- 60° in 3° increments with a total dose of 100 e^-^/Å^2^ using a dose symmetric scheme. Each tilt was collected as a movie with 3 subframes, and an exposure of 0.36s per tilt. Tilt series were collected at a defocus of – 4 µm. Motion correction and restacking of the tilt series was performed using MotionCorr2 using scripts written in house [21]. Individual tilt images were then [22]denoised using TOPAZ and tilt-series were imported in IMOD 4.12.11, binned 4-8x and tomograms were reconstructed using standard reconstruction workflows using the ETomo suite [23]. Missing wedge correction was performed using Isonet [24]. and final visualization of the reconstructed volumes was done in 3dmod and ChimeraX [25].

## RESULTS

### Conventional negative stain TEM introduces artifacts and occludes some native details

Using a soil sample from a managed agricultural field site in Prosser, WA which was enriched for viruses using an approached optimized for downstream genomics analysis, we performed Negative Stain TEM to get an initial image of the sample. While the typical fields of view for Negative Stain TEM can allow researchers to pick and choose the best local area and particle of interest, we chose random locations to provide more reproducible examples of best imaging conditions. As seen in Figure S1, two random locations within the same ∼50 micron x 50 micron grid square show vastly different backgrounds. While similar elongated symmetrical particles with intact tails can be identified in both images with the arrowheads, the two particles show a putatively intact virus with genomic content still inside the capsid (red arrowhead) and a particle that clearly has stain inside the capsid indicating the DNA/RNA content was already released or the particle itself is damaged and not intact (white arrowhead). Other particle morphologies can be seen but the images also show aggregation, stain artifacts and particle spatial overlap. For example, while a bottle-shaped phage is seen in **Figure S1A** (white asterisk) the stain prevented observing any high-resolution features like surface glycoproteins which are expected to be present on that morphotype[26, 27].

### Filtration improves cryo-EM imaging of viruses

Suspensions of native viruses isolated from soil generally have low abundance compared to the amount of material needed for protein crystallography and they contain nanometer to micron scale debris which complicate cryo-EM. Debris that is usually not a problem in conventional negative stain electron microscopy or multi-omics becomes a hindrance in cryo-EM for observing any viruses or virus like particles (VLPs) as the largest structures tend to set the local ice thickness gradients (**Figure 1, grid *1**).

**Figure 1.**
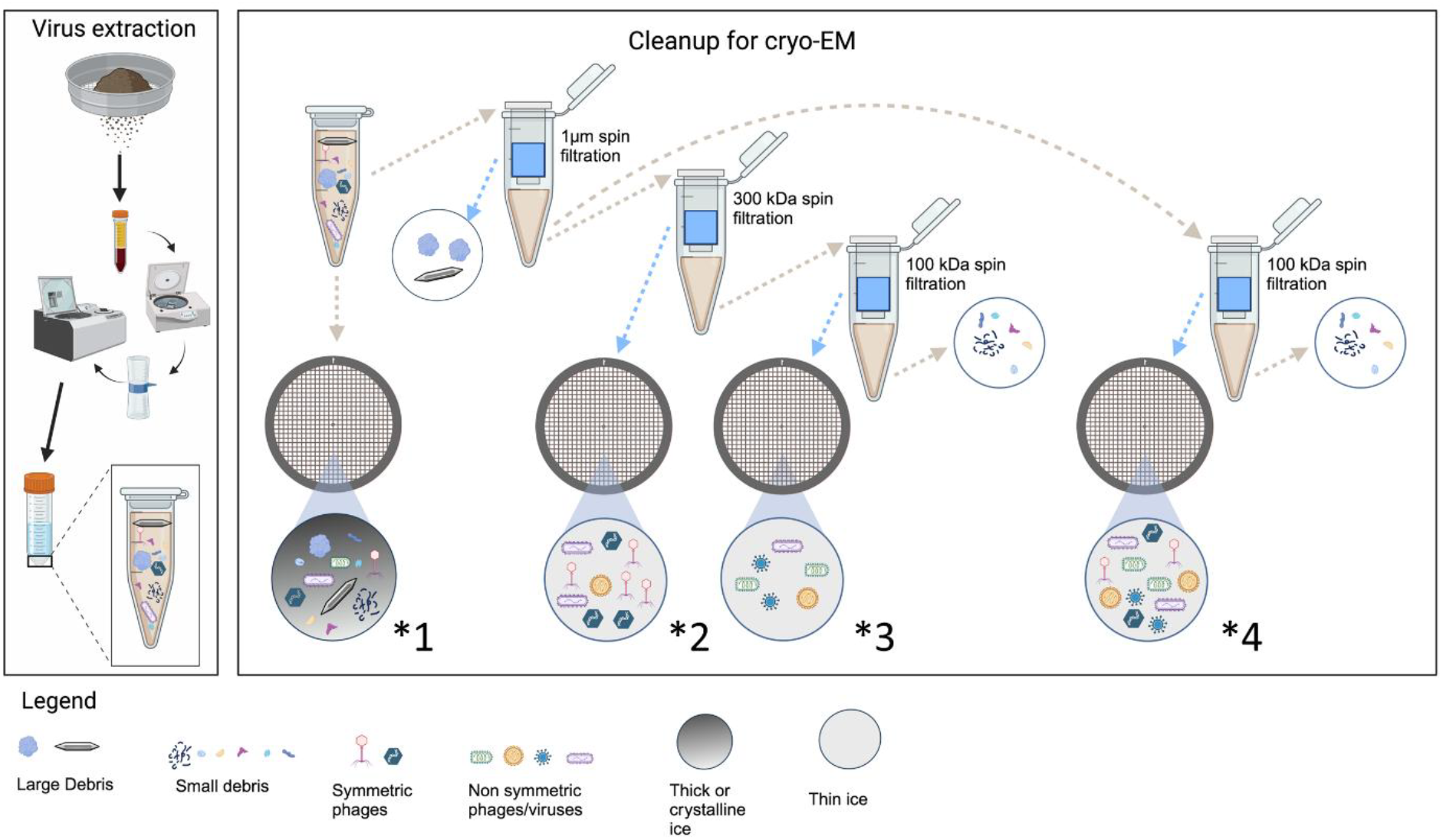
Workflow cartoon of methods to prepare soil samples for cryo-EM. Virus extraction - Soil samples were processed per Santos-Medellin et al to extract viruses. Clean-up for Cryo-EM – Suspensions of viruses natively isolated from soil contain viruses as well as nanometer to micrometer size debris. Soil suspensions frozen on grids as-is and imaged with cryo-EM contained lots of debris and thick ice which hindered visualization of virus particles (*1). However, soil suspensions passed through a 1 µm pore size Pleuriselect filter got rid of larger debris in the retentate and allowed the filtrate containing the viruses and smaller debris to be subsequently further concentrated and fractionated using 2 separate approaches. In the first approach, sequential filtering steps help separate different size viruses. Here the filtrate from the Pleuriselect filter can be concentrated through a 300 kDa MWCO filter to concentrate viruses and get rid of nanometer sized debris. This sample resulted in thin ice and facilitated visualization of viruses. Cryo-EM images collected on the retentate (*2) capture most of the virus diversity. Filtrate from the 300 kDa MWCO filter can be further processed through a 100 kDa MWCO filter. Imaging the retentate (*3) showed fewer total viruses that were mostly smaller virus like particles, while imaging the filtrate showed mostly junk protein. If desired the sequential approach allows for visualizing the differently sized viruses. Alternatively filtrate from the Pleuriselect membrane can be directly loaded on a 100 kDa MWCO filter. The retentate (*4) from this approach captures the entire mixture of small and large viruses while and most small protein debris is removed to provide quality thin ice to directly visualize the viral diversity. (Created using BioRender.com).

Using direct freezing of the as-delivered initial suspension, we observed thick ice and the presence of large aggregates and mineral debris despite the original preparation for viral enrichment including filtration through a 0.22 µm pore size membrane filter (**Figure 2** and **S2**). We therefore employed a cleanup step to get rid of debris and aggregates that might occlude or hinder visualization of viruses and VLPs in cryo-EM imaging. In principle, cryo-EM imaging with a phase plate and energy filter can image specimens up to 1 micron in thickness. Our cleanup step therefore consisted of centrifuging the suspension through a 1 µm pore size membrane Pleuriselect filter to separate out the larger micron-sized debris. We then subject the cleaned filtrate to simultaneous concentration and size selection filtration using 300 or 100 kDa MWCO spin columns to further improve the sampling efficiency and image quality.

**Figure 2.**
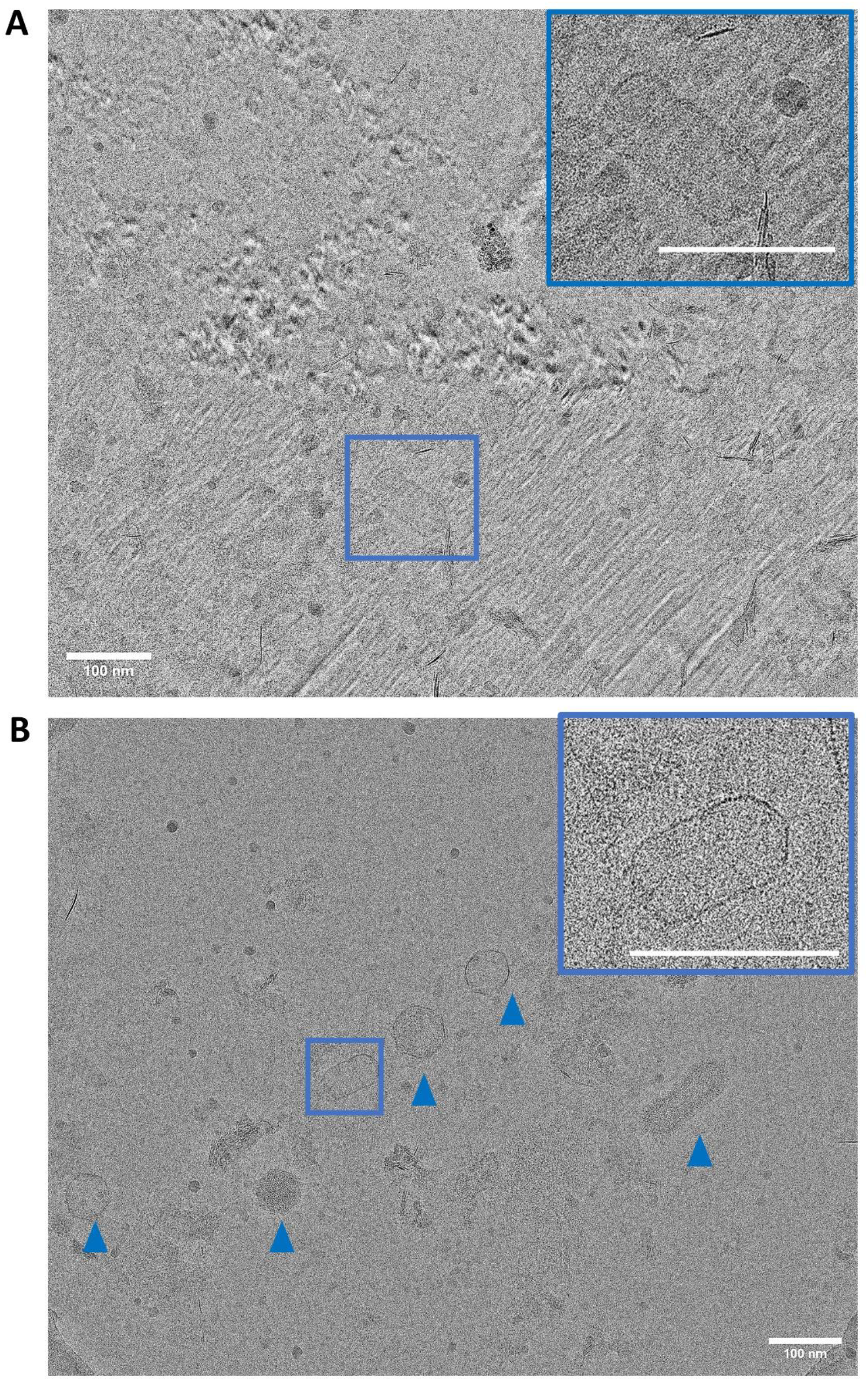
Ice quality comparison of grids frozen before and after the cleanup step. (A) A cryo-EM image at 42000 x nominal magnification showing overall devitrified crystalline ice. The inset is a zoomed in view of a phage (blue boundary) which shows the crystalline ice hindering visualization of the phage morphology. (B) A cryo-EM image from the sample post cleanup which shows vitreous ice that allows visualization of multiple phage and virus particles (blue triangles). The inset is zoomed in view of a phage in blue boundary showing glycoprotein spikes on an unidentified elongated virus. Scale bar = 100 nm

Following this simple cleanup, we observed that at lower magnifications (135x-580x) the atlases and montages appeared roughly similar between the as-delivered and final cleaned up samples **(Figure S2 A-C, S3A-C)**. However, images and montages collected at 11,500 x **(Figure S2D, S3D)** showed clear differences where the as-delivered sample depicted high amounts of debris which was absent from the cleaned samples **(Figure S2E, S3E)**.

Any protein macromolecules like viruses frozen for cryo-EM should be ideally present in a layer of uniform vitreous ice only slightly thicker than the target macromolecule to maximize contrast and resolution. To compare ice quality on the grids with virus suspensions frozen as-delivered and after filtration, we collected images at 42,000x where the presence of ice crystallization artifacts from uneven vitrification would be obvious. Crystalline ice has unit cell with dimensions around 3.4 Angstroms which results in prominent spots visible in the Fast Fourier Transform of the image or in severe cases the lattice is visible in the direct image. Due to this, images which have crystalline ice cannot be used for high resolution structural studies. As expected, images of the as-delivered virus suspension showed aggregates, debris as well as devitrified ice **(Figure 2A)** with multiple microdomains showing tell-tale signs of crystalline ice. While a bacilliform phage like particle was visible in the field of view, no high-resolution features could be clearly seen even in magnified views **(Figure 2A inset)**. However, after filtration and removal of the larger debris, the cryo-EM image (**Figure 2B**) of the virus suspension showed uniform vitrified ice (confirmed by absence of any detectable lattice in the direct image nor any diffraction spots in the FFT of the image, data not shown). The vitreous ice, with minimal debris from particle aggregation, allowed visualization of several phages and one elongated virus in the field of view. A zoomed in view of the virus particle **(Figure 2B, inset)** showed similar morphology as that in Figure 2A but here the vitreous ice allowed better visualization of the capsid and other features.

To estimate how many viruses may have escaped observation in the 300 kDa MWCO retentate, we further concentrated the 300 kDa MWCO filtrate over a 100 kDa MWCO spin concentrator and imaged the resulting retentate (Figure 1, *3). Unsurprisingly, we mostly observed vlps with diameters between 20-50 nm and an occasional virus. By quantifying the number of observed phages seen in roughly equivalent datasets of around 1100 captured micrographs from each of the 300 kDa MWCO and 100 kDa MWCO retentates, we concluded that the 300 kDa spin concentration step is efficient to capture most (>90%) of the viral diversity from our current sample and the 100 kDa spin concentration post 300 kDa spin concentration may not be absolutely necessary beyond getting rid of small debris that can obscure high-resolution imaging of smaller viruses. Thus, if the main goal is to evaluate the diversity of the viromes, a combination of Pleuriselect 1 micrometer filtration followed directly by 100 kDa MWCO filtration (Figure 1, *4) provides the best option for capturing all viruses and vlps present in solution while removing both large and small debris that hinder high-quality imaging.

### Direct visualization of viral diversity in soil extracts

While previous negative stain TEM studies [28-32] of soil extracts and high-resolution cryo-EM structural characterization of purified viruses have provided detailed classification of symmetric viruses, we decided to classify any detected viruses based on known and unknown morphotypes to fully capture the diversity. The most readily identifiable morphology consisted of tail-less polyhedral phages and phages with tails of varying length **(Figure 3A-B)**. Figure 3B.1 shows a bacilliform phage while 3B.2 shows a myovirus-like phage with relatively thicker tail, and Figure 3B.3 a podovirus-like phage with a very short tail. Figure 3B.5 shows a siphovirus-like phage where the head, tail and tail fibers are clearly visible. Our workflow also clearly enables visualization of features like contractile and non-contractile tails **(Figure 3B.2, 3B.1,5)** as well as glycoproteins on several symmetrical phages **(Figure 3B.4)**. Diameters for the icosahedral phage heads ranged from 50-80 nm whereas the bacilliform phages had 100-120 nm long heads with 40 nm diameter. Several images showed the presence of long thin tube-like structures with 10-20 nm diameter and of varying lengths (200 nm – 1 µm) which could not be visually assigned to any group of viruses **(Figure 3C.1,2, Figure S4 C)** and could be remnants of [31]broken tails or contractile sheaths or something else currently undetermined. Another major morphotype observed had an overall round shape, diameter <50 nm and no visible glycoproteins on the surfaces **(Figure 3C.3,4,5, Figure S5 C)**.

**Figure 3.**
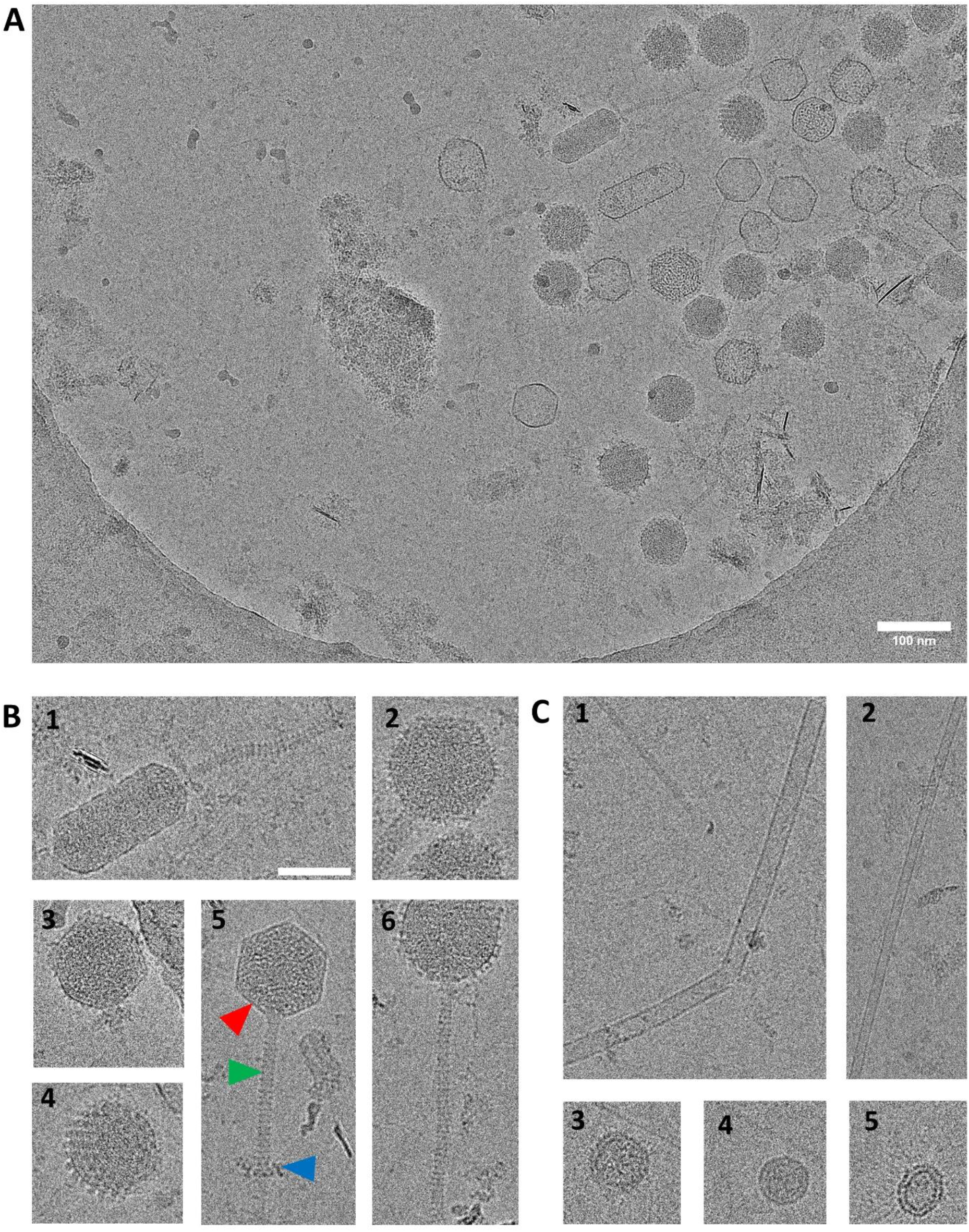
Cryo-EM images of symmetric viruses in filtrate of the sample 100 kDa MWCO filter. **A)** Cryo-EM image showing multiple phages with symmetric shapes. Tailed and tail-less phages along with other icosahedral viruses. Phages display both icosahedral and bullet shaped heads. B) Zoomed in view of 3 select phages from (A) shown. 1. A bacilliform phage showing a long non contractile tail and siphovirus like morphology. 2. Phage showing a thick contractile tail having myovirus like morphology. 3. Podovirus like phage showing an extremely short tail (green triangle) (Phages in 1,2,4 are zoomed in views of phages from 3A). 4) Tail-less phage with glycoproteins visible on the virus surface. 5. Phage showing intact head (red triangle), contractile tail (green triangle) as well as tail fibers (blue triangle). 6. Phage showing a slightly rounded head relative to the phage shown in 3A.6. C) 1 and 2 are tube like VLPs of differing diameters abundantly observed in images. 3, 4, and 5 are VLPs which are <50 nm in diameter and have no visible glycoproteins on the surface. Scale bar = 50 nm for B and C.

In addition to the tailed and polyhedral viruses which are readily seen with negative stain TEM and above with the cryo-EM workflow described here, we also identified several virus morphotypes which have only be occasionally reported [28] and some novel ones which have never been directly visualized before. These novel morphotypes could belong to any of the multiple genera reported from soil viromes which currently have no morphology associated with them. These virus morphotypes were broadly divided into groups based on either shape, size or other visual similarities **(Figure 4)**. The presence of an intact membrane and visible glycoprotein spikes of any shape and size were considered as pre-requisites for a visual “feature” observed in an image to be counted as a virus. Round viruses with diameters between 50-200 nm and visually different surface glycoprotein spikes made up the numerically second largest group (**Figure 4.1)**. A diverse group of morphotypes showed the presence of a very thick coat surrounding the viral capsid/core **(Figure 4.2-6)**. The sizes of these viruses ranged from 20-157 nm with coat thickness from 15-35 nm. We observed an inovirus like morphotype (caterpillar like) that had a cylindrical central core dotted with putative glycoprotein spikes orthogonal to the axial virus core **(Figure 4.7,8)**. Bottle shaped viruses with diverse morphologies **(Figure 4.9, 4.10**) and filamentous viruses **(Figure 4.11)** were also occasionally observed. Interestingly, the images of bottle shaped viruses by cryo-EM clearly show a glycoprotein-like coat that is not visualized in a similar negative stain image (compare Figure 4.10 with Figure S1A white asterisk). Another group of viruses with diameters ranging from 30 to 85 nm showed the presence of fiber like glycoprotein like projections between 31-47 nm in length (here called LG viruses) **(Figure 4.12)**. The LG viruses are visually similar to morphotypes described by Fischer *et al* as viruses with capsid fibers[14]. Of note, the size/dimension of virus morphotypes did not have a linear correlation with either the measured thickness of the external coat or length of the glycoprotein projections. Some polymorphic viruses with irregular shapes were also observed which have not been reported before in literature **(Figure 4.13,14)**. While Figures 3 and 4 document the library of 2D morphotypes identified by this imaging approach, we also summarize the results in Figure 5 to illustrate the diversity more graphically.

**Figure 4.**
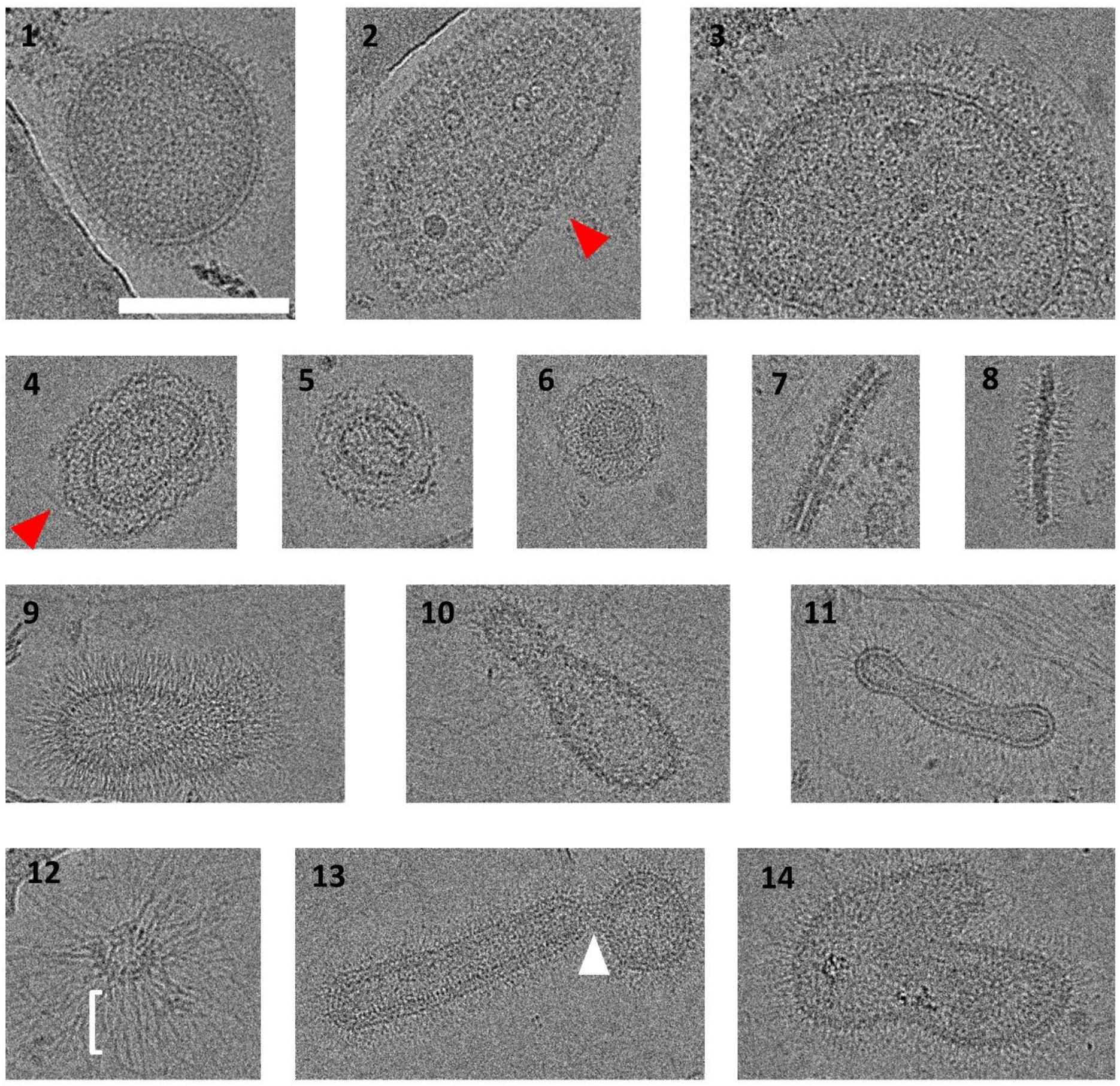
Cryo-EM images of polymorphic viruses in filtrate of the sample 100 kDa MWCO filter. 1) Example of a round phage. 2-6) Diverse uncharacterized phages displaying a wide range of diameters and a thick outer coating/layer external to the virus core (red triangle). 7-8) Caterpillar like appearance of a virus with long thin central core and glycoproteins on the surface putatively belonging to the Inoviridae Family. 9-10) Tube shaped viruses with diverse morphologies 11) Elongated or filamentous phage. 12) Phage showing unique long projections which could be glycoproteins or glycans (LG, red arrows) with the white parenthesis indicating the length of the glycoproteins. 12) A polymorphic vlp showing a head and body morphology on either side of a narrow neck (white triangle) 13) Polymorphic phage having plasmavirus like irregular morphology. Scale bar 100 nm.

**Figure 5.**
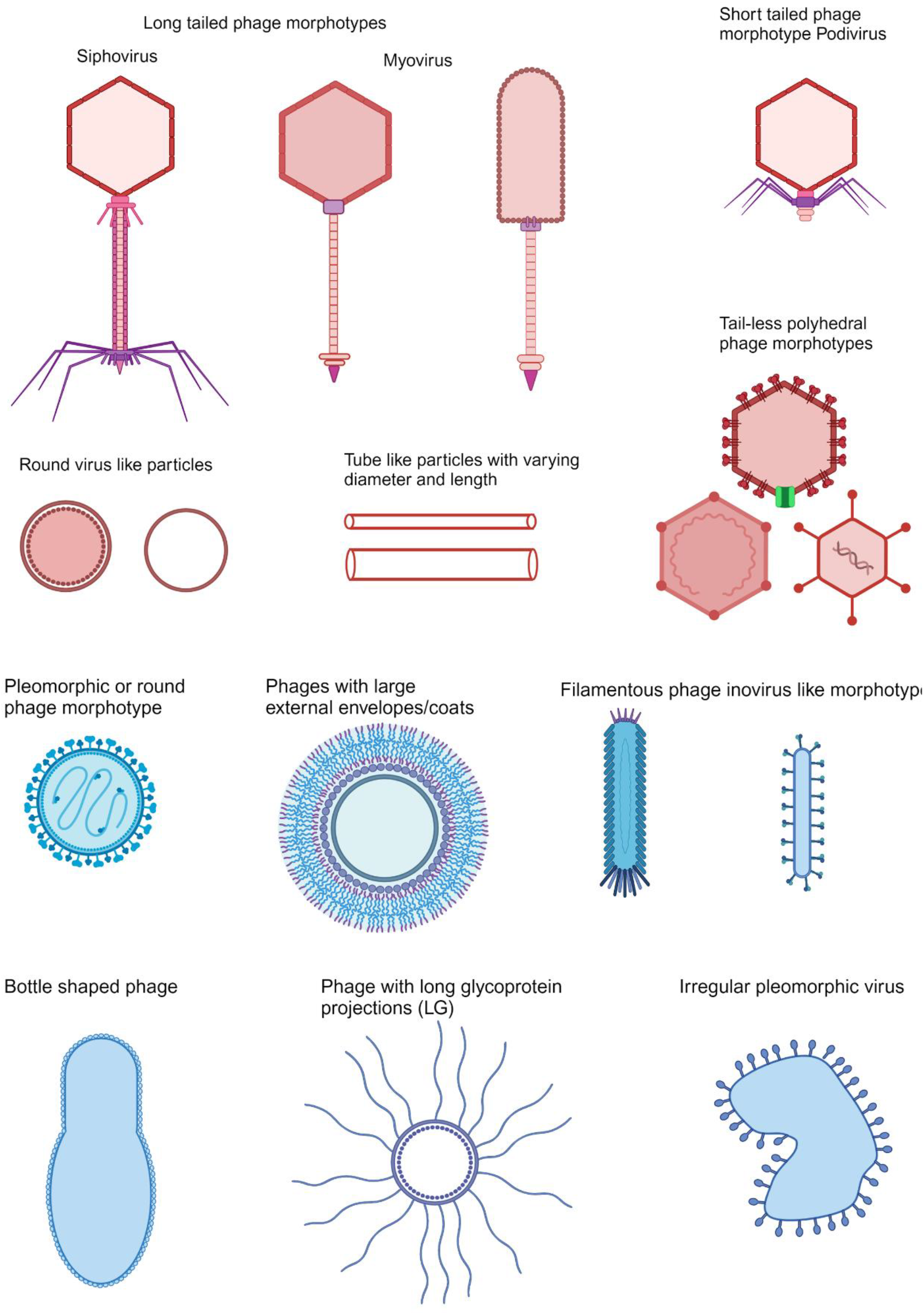
Cartoon representation of virus morphotypes in current study. The brown/red morphotypes have been reported in previous studies and were also observed in the current cryo-EM analyses. The blue morphotypes are mostly uncharacterized, diverse phages some of which have not yet been reported in literature. (Created using Biorender.com)

One of the main benefits of cryo-EM, besides imaging samples in near-native conditions without crystallization, is that it can be used to both produce a 2D library of images as above and obtain 3D volumetric information using tomography. Since the polymorphic viruses have not been widely studied and their overall shape can be hard to recognize in 3D from a single 2D projection, we performed cryo-electron tomography on a small-subset of the viruses **(Figures S4 and S5)**. This approach allows tracing the contiguous and intact membranes for the polymorphic phages as well as detecting glycoprotein extensions and in some cases density inside the viruses that could correspond to packed DNA/RNA.

## Discussion

Negative staining and conventional TEM has been widely used to visualize viruses and VLPs from soil environments, but a number of technical and operational problems have plagued this technique and hindered viral research. While negative stained TEM is a quick and easy way to visualize viruses, a recent report delved into the gradual decline in quality of electron microscopy images seen over the years by evaluating hundreds of publications where the loss was largely ascribed to a loss of expertise and training [33]. Additionally, previous methods for extracting viruses from soil have sometimes been too harsh or biased towards tailed phages [34]. Most studies of TEM imaging of tailed viruses isolated from soil have reported broken tails or loss of tails, where the frequency of loss of tail is inversely proportional to the tail length [6, 35]. Here we have demonstrated an alternate technique for visualizing viruses from challenging samples that employs highly selective but gentle filtration and cryo-EM to mitigate or eliminate many of the problems that plague negative stain TEM images.

So far, cryo-EM of soil viruses has been restricted to studies where a single virus isolated from soil sample can be propagated and purified in quantities amenable for cryo-EM single particle studies [36]. Here we overcome this challenge/bottleneck and demonstrate the ability to visualize the native viral community isolated directly from soil using cryo-EM by using common and simple physical separation techniques. We used microcentrifuge spin columns to simultaneously remove unwanted debris and to concentrate the sample to increase likelihood of detection on the grid which overcame two of the main bottlenecks. The resulting grids were then imaged using semi-automated cryo-EM screening and data collection where we able to detect at least one virus or VLP in ∼70% of the images collected on the 300 kDa MWCO retentate. Since most of the virus suspension loaded on a grid gets blotted away prior to vitrification, the virus library we have obtained **(Figure 5)** cannot yet be truly representative and it is possible that several morphotypes could not be imaged. However, this drawback also extends to libraries obtained using negative stained TEM and our work sets the foundation for being able to visualize the viral diversity with cryo-EM while future work can focus on optimizing the loading and sampling efficiency to capture more of the diversity. For example, multiblot methods are known to increase the local concentration of particles trapped in cryo-EM grid holes even when using wicking, while lossless sample deposition and vitrification can be accomplished using the Vitrojet [37] and other similar preparative instruments.

Another way to validate the diversity is linking the observed cryo-EM 2D and 3D data with omics analysis on the same samples. While beyond the focus of the current study, orthogonal multi-omics would be needed to provide more information about the morphotypes imaged in this study to confirm that a novel virus morphotype is truly an infectious virus. For example, gene transfer agents (GTAs) appear and function like phages but carry fragments of cellular genome resulting in horizontal gene transfer. It would be very challenging to distinguish between a bacteriophage and a GTA purely based on 2D images [38]. Additionally, some of the round virus morphotypes we have observed may be a result of cellular blebbing. Without further orthogonal studies, it is not possible to unambiguously assign all phage morphotypes to the Class *Caudoviricetes* [17].

Given estimates of the diversity of viruses in soil, there is ample opportunity for this work to be replicated on soil samples from numerous diverse locations, during different times of the year to obtain a broader representation of viruses resident in soil. Further, for higher resolution studies, soil virus extractions could be processed using the virus reduction approaches used in aquatic virology to selectively propagate a few viruses in high copy number [39, 40]. Complete purification and isolation of these viruses would not be needed; their selective amplification would provide suitable samples for high-resolution cryo-EM/ET studies using sub-tomogram averaging, while also allowing verification of host-virus infectivity. Hypothetically, a dataset with just a few hundred symmetric virus particles could be used to obtain 2D classes and sub-nanometer resolution 3D maps. The selectively amplified viruses could be used to infect target host cells and samples can be frozen at various stages of infection to perform cryo-ET, which will provide structural details of interaction of host-virus systems from soil in unprecedented resolution.

## Acknowledgement and Funding

The research was performed using EMSL (grid.436923.9), a DOE Office of Science User Facility sponsored by the Biological and Environmental Research program located at PNNL. A portion of this research was performed on project award (10.46936/fics.proj.2022.60449/60008585) under the FICUS program and used resources at the DOE Joint Genome Institute and the Environmental Molecular Sciences Laboratory, which are DOE Office of Science User Facilities. Both facilities are sponsored by the Biological and Environmental Research program and operated under Contract Nos. DE-AC02-05CH11231 (JGI) and DE-AC05-76RL01830 (EMSL). A portion of this research was supported by the DOE Office of Biological and Environmental Research FWP 74915 and D7G33.

## Author Contributions

JEE, ADP conceived this project. ADP, RWM, TA, AEZ performed all the experiments and ADP performed all the data analyses. JEE, WCN and KSH provided supervision and research support (funding). All the authors contributed to drafting and editing the manuscript.

## Supplementary Figures

**Figure S1.**
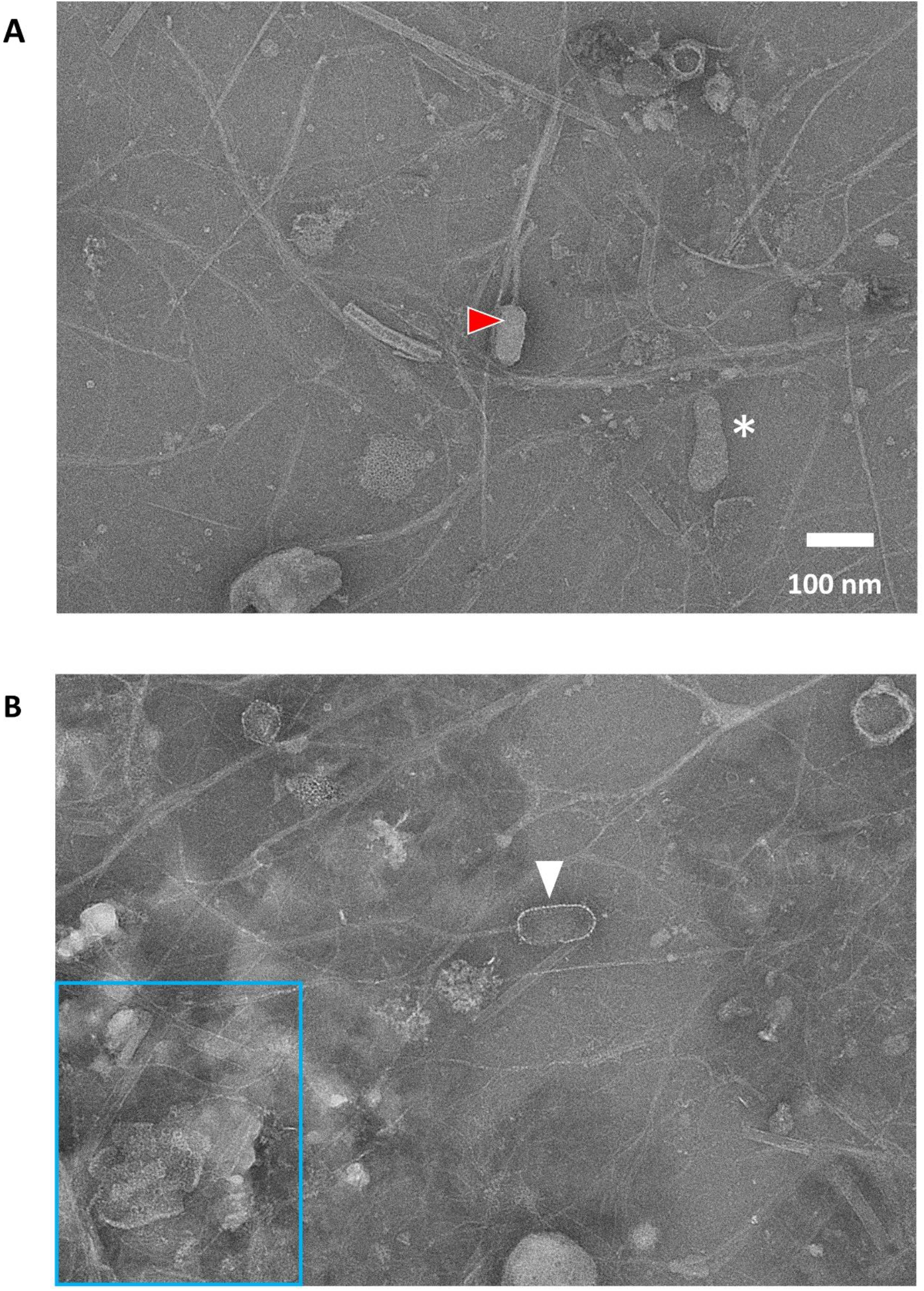
Negative stained image of viruses natively isolated after the Pleuriselect step. Conventional negatively stained images of viruses collected at 42000 x nominal magnification. A) Example of a good negative staining where only the background has been stained and phage shape is clearly visible (white triangle) B) Example of an overstained image where stain seems to have penetrated inside the phage head (read triangle). The images show a lot of background junk and loss of high-resolution features. Area in blue boundary shows aggregates in the field of view which prevents identifying any VLPs due to spatial overlap.

**Figure S2.**
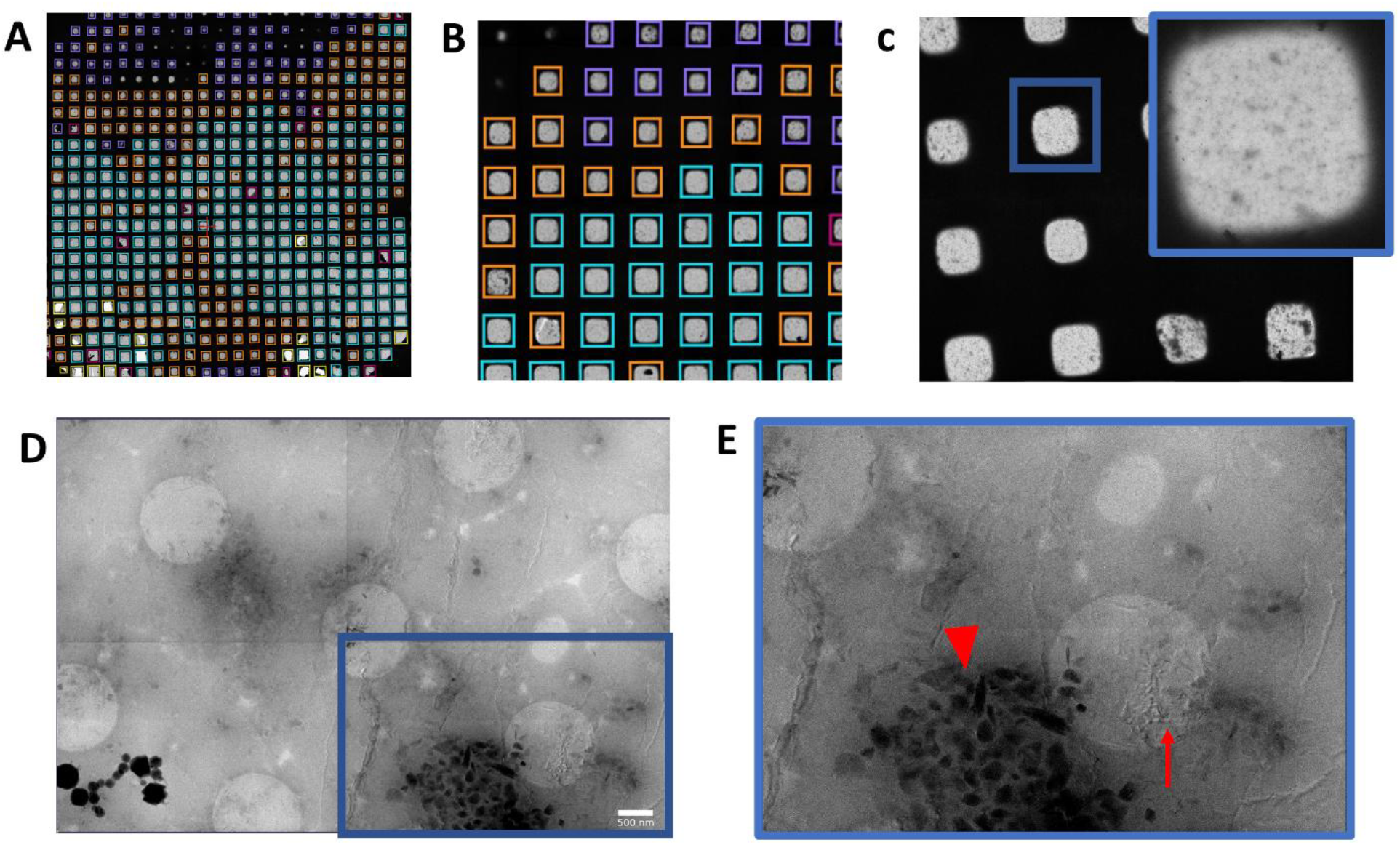
Comparison of scales of imaging for virus samples before purification. A) A stitched atlas images collected at very low magnification showing the entire cryo-EM grid showing. Squares on the grid are colored from relatively thick (purple) to thin (blue) with the blue colored squares having ice thickness ideal for cryo-EM imaging to B) A zoomed in area of the atlas level magnification shows no visible difference in the grid with samples before or after purification. C) A higher mag view of the grid with an individual square (blue border and inset) that shows no visible difference in ice quality. D) A 2x2 montage of individual images collected at 11500x magnification for the sample before Pleuriselect cleanup. At this magnification, the ice begins to show irregularities. E) Zoomed in image of the individual image in the montage shows presence of debris (red triangle) as well as crystalline ice (red arrow). Scale bar for D and I – 500 nm. Each circular hole is 1 µm in diameter.

**Figure S3.**
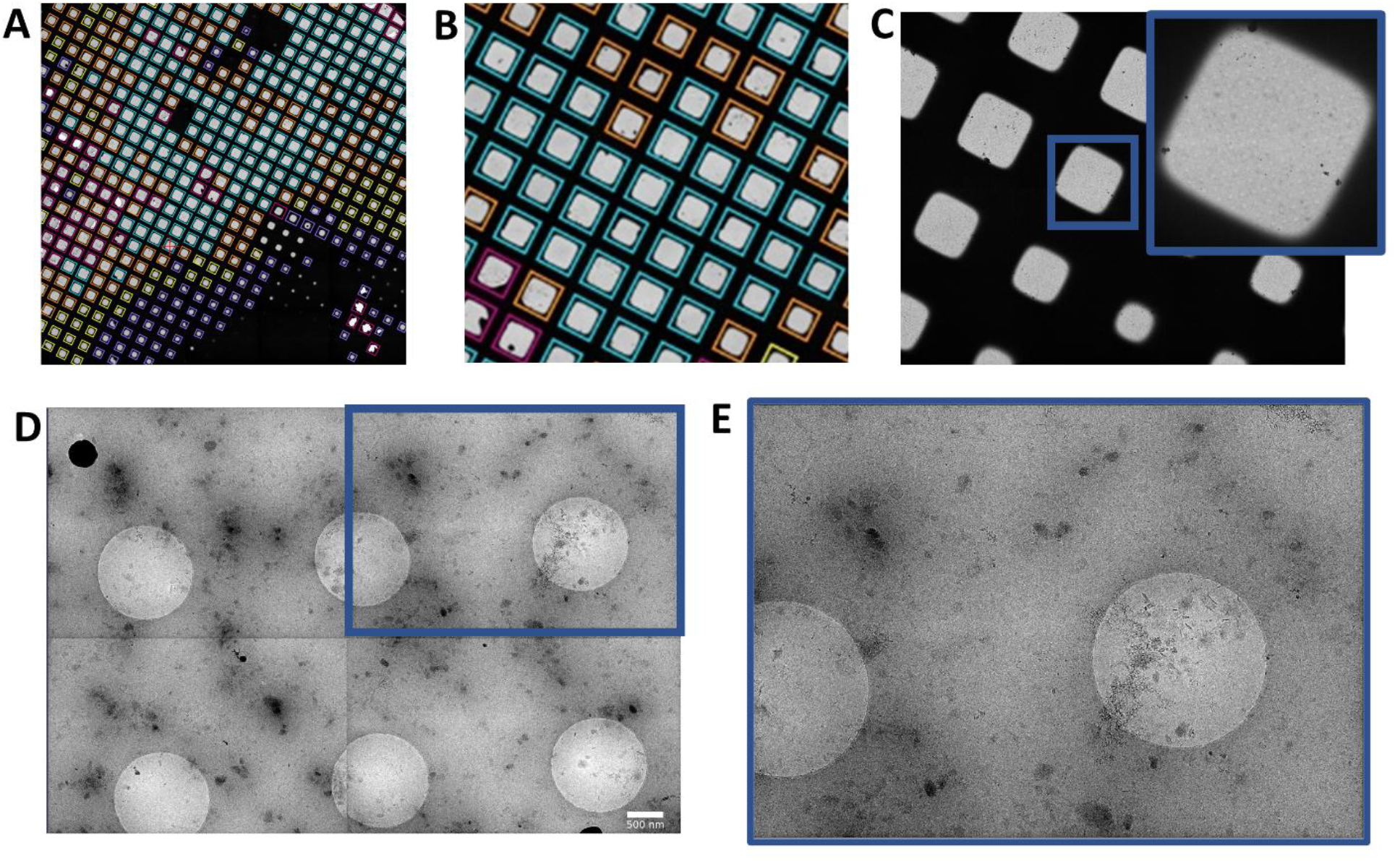
Comparison of scales of imaging for virus samples after purification. A) stitched atlas images collected at very low magnification showing the entire cryo-EM grid showing. Squares on the grid are colored from relatively thick (purple) to thin (blue) with the blue colored squares having ice thickness ideal for cryo-EM imaging to B) A zoomed in area of the atlas level magnification shows no visible difference in the grid with samples before or after purification. C) A higher mag view of the grid with an individual square (blue border and inset) that shows no visible difference in ice quality. D) A 2x2 montage of individual images collected at 11500x magnification for the sample after Pleuriselect cleanup. E) Zoomed in image of the individual image in the montage shows no detectable. No visible large debris or crystalline ice seen in the montage or the zoomed in image. Scale bar for D and I – 500 nm. Each circular hole is 1 µm in diameter.

**Figure S4.**
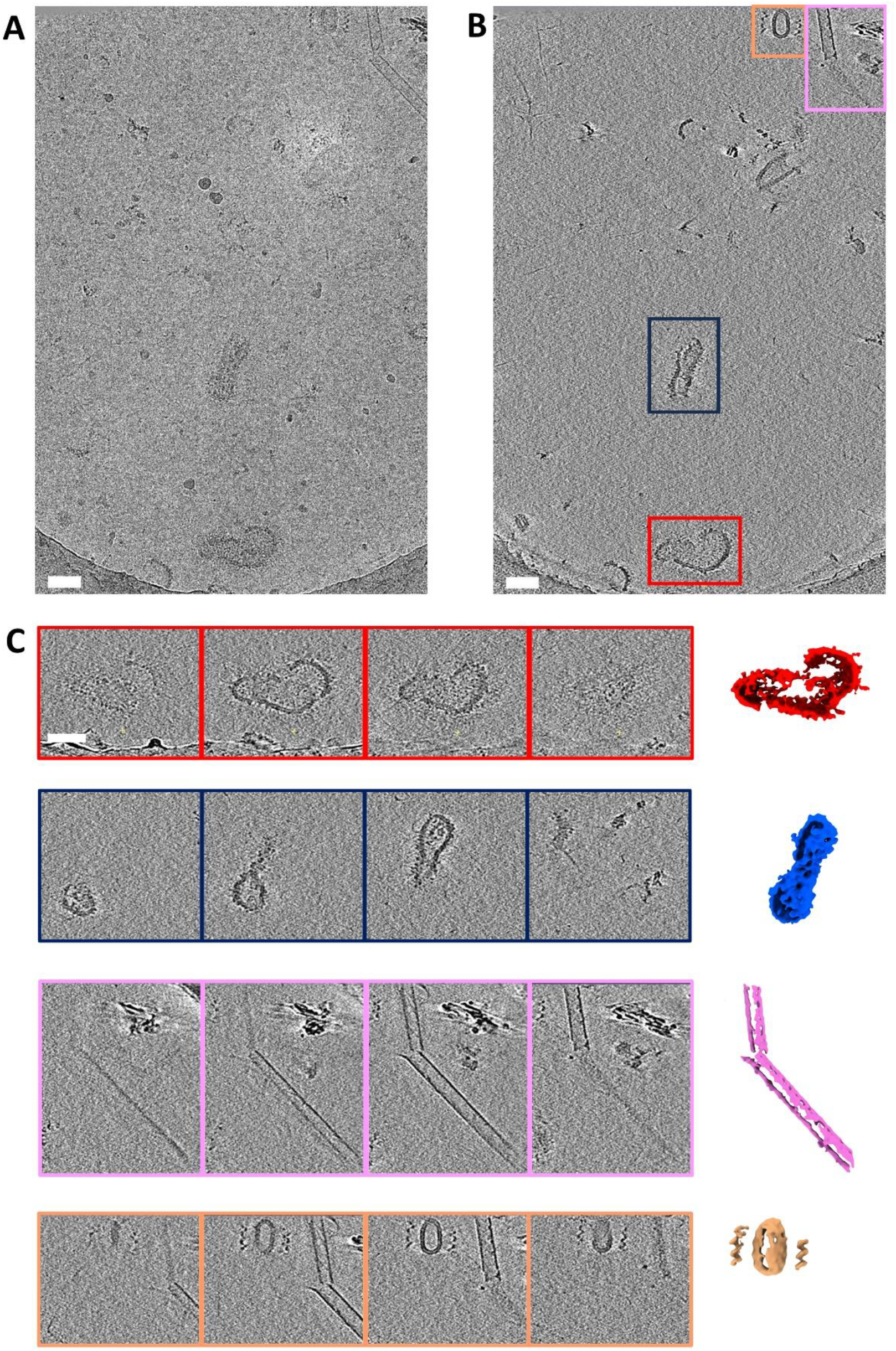
Cryo-tomographic imaging of viruses natively isolated from soil. A)Zero-degree tilt image of the tomogram with diverse morphotypes. B) Slice through the reconstructed tomogram with 3 morphotypes in in the red, blue and yellow boundary. C) Successive Z slices through the tomogram for each of the individual morphotypes tracing appearance and disappearance of the viruses/vlps. False color rendering of the virus/vlps are shown as visualized using ChimeraX. Red – a comma or bean shaped vlp showing intact membranes and glycoprotein spikes visible on its surface. Blue – An elongated virus morphotype with blebs internal to its membrane and glycoprotein spikes on the surface. Pink – A tube like morphotype observed with uncharacterized material internal to the tube. Beige – Morphotype with a thick coat surrounding the central core. Slices in pink and beige boundaries have overlapping regions. All scale bars = 25 nm

**Figure S5.**
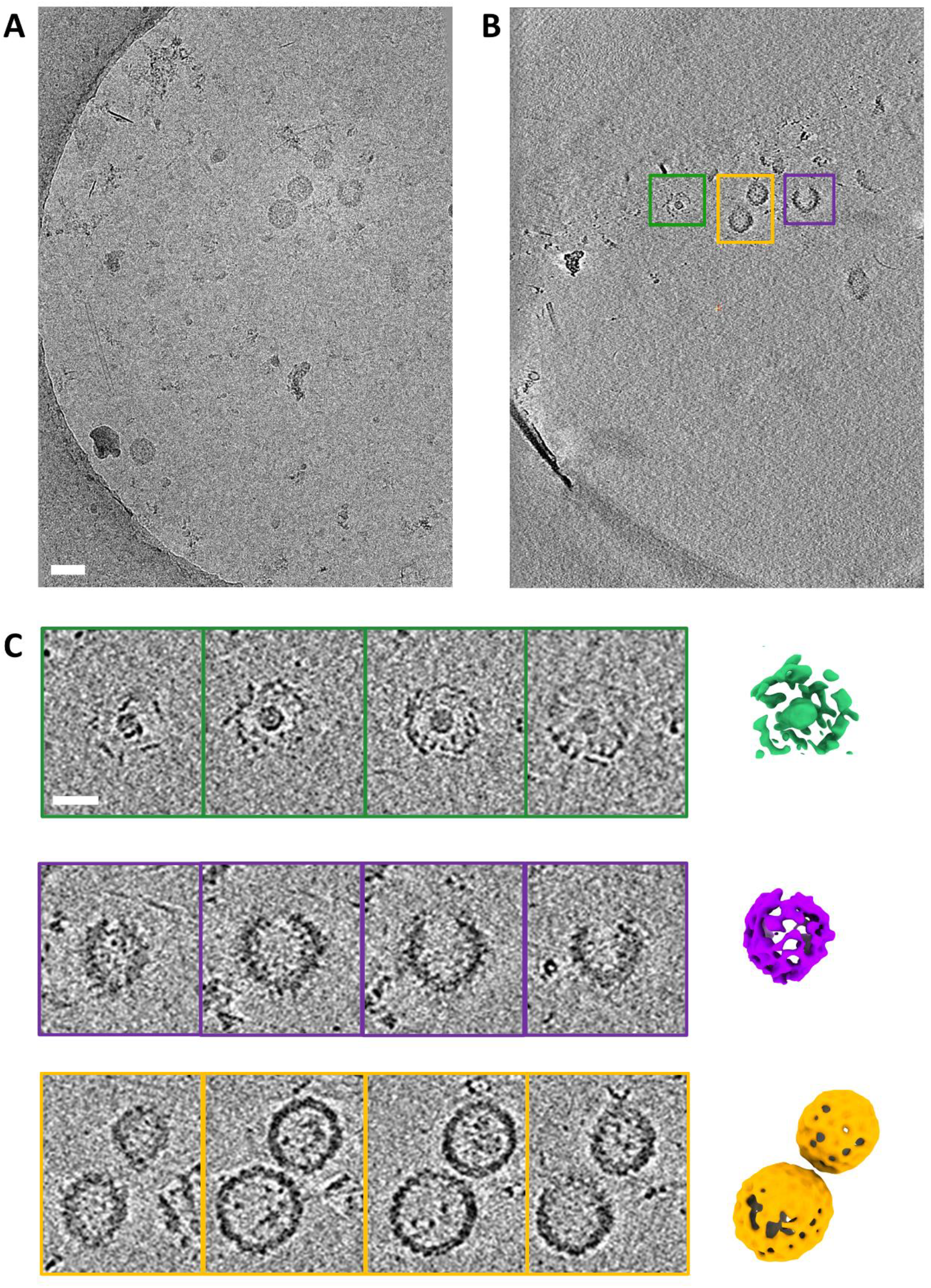
Cryo-tomographic imaging of vlps natively isolated from soil. . A)Zero-degree tilt image of a tomogram with diverse morphotypes/vlps B) Slice through the reconstructed tomogram with 3 morphotypes in in the green, violet and gold. C) Successive Z slices through the tomogram for each of the individual morphotypes tracing appearance and disappearance of the viruses/vlps. False color rendering of the virus/vlps are shown as visualized using ChimeraX. Green – A round morphotype with a small core and a thick coat. Violet– A curved morphotype. Gold – Two round vlps very close to each other. No glycoproteins visible on their surface. All scale bars = 50 nm

